# A new method for the reproducible development of aptamers (Neomers)

**DOI:** 10.1101/2023.12.19.572437

**Authors:** Cathal Meehan, Erika L. Hamilton, Chloé G. Mansour, Soizic Lecocq, Cole J. Drake, Yisi An, Emily Rodrigues, Gregory Penner

## Abstract

The development of aptamers has been almost exclusively performed based on the SELEX method since their inception. While this method represents a powerful means of harnessing the in vitro evolution of sequences that bind to a given target, there are significant constraints in the design. The most significant constraint has been the reliance on counter selection on off-targets to drive specificity. Counter selection has not been as effective at driving aptamer specificity as the presence of immune tolerance, the capacity of the immune system to remove antibodies that bind to host targets, is for antibody development. This deficiency has constrained the commercial efficacy of aptamers to date. These limitations have been addressed with our design of a novel platform for aptamer identification. This new design is based on what we refer to as a Neomer library with sixteen random nucleotides interspersed with fixed sequences. The fixed sequences are designed to minimize the potential for hybridization, such that secondary structure is driven by the random nucleotides. The use of sixteen random nucleotides reduces the possible library sequences to 4.29 × 10^9^. This enables the application of the same sequences to either the same target or different targets while maintaining a high level of structural diversity. In effect, this introduces the capacity for reproducibility in aptamer selection and in silico approach to replicating immune tolerance We provide here an overview of the new method and a description of the performance of aptamers selected for interleukin 6 developed using this approach.

## Introduction

The invention of aptamers by four different groups between 1989-1990^1–4^ introduced the principle of Systematic Evolution of Ligands by EXponential enrichment (SELEX) for the enrichment of DNA or RNA oligonucleotide sequences from random libraries for specific targets. Since then, there has been continued improvement of the SELEX process to improve aptamer specificity and affinity. In 1992 Ellington et al. introduced negative SELEX with counter selection against the solid support used to immobilize the target^5^ and in 1994, Jenison et al. added counter selection against similar targets to enhance specificity^6^. SELEX methodology has also been applied in different platforms including capillary electrophoresis (CE) SELEX^7–9^, which increases the stringency of selection; microfluidic SELEX^10,11^, which reduces the amount of target required for selections; and Cell SELEX^12^, which allows for selection of targets in their native environment and agnostic selection against cell types. These adaptations of SELEX provide many advantages, but they are still constrained by their core reliance on the SELEX concept.

The SELEX procedure consistently has a library design featuring a contiguous random region flanked by two polymerase chain reaction (PCR) primer recognition sequences. The size of the random region varies from 25 to 80 nucleotides (nt) and depends on different factors, such as the size of the target^13^. Research indicates that the length of the random sequence directly influences the structural diversity of the SELEX library, with smaller random regions yielding less structural diversity^14^. A minimum number of homologous nucleotides is necessary to form stable hybridizations, and the hybridization of at least three nucleotides between different regions of the aptamer are required for the formation of hairpin loops^15^.

To optimize structural diversity, it has become common practice to design libraries consisting of 40 random nucleotides. A random region of 40 random nucleotides consists of 1.2 × 10^24^ possible sequences. Because it is impractical to work with this many sequences in a selection process, selections generally start with an initial naÏve library containing 1 × 10^15^ random sequences. This subset represents a proportion of 8.27E-10 of the total sequence solution space. Such a small subset of the total sequence solution space creates two constraints. First, there is an average of one copy for each unique sequence in the initial naÏve library before the first round of selection. This means that there is a possibility that sequences that would show binding to the target aren’t identified due to stochastic variation in the binding solution space. This also means that multiple iterative rounds of selection are needed for sequences to sufficiently enrich for adequate characterization. Prior to each subsequent selection round, it is essential to amplify the selected library which introduces additional technical variation due to PCR bias^16^ into the selection process.

Secondly, the small subset of the total sequence solution space means that selection is not reproducible. It is not possible to start selection with the same subset of initial naÏve sequences. Lack of reproducibility between selections is also a constraint with antibody production as the pre-infection repertoire of antibodies show large variation on an individual basis^17^.

A key biological difference between antibody development and aptamer development is the presence of immune tolerance within the antibody development system. Immune tolerance is the capacity of the immune response system to remove antibodies that bind to pre-existing host epitopes, thus preventing the immune system from damaging the host^18^. The in-vitro nature of aptamer development means that this step is not present. Moreover, the lack of reproducibility implicit in SELEX means that it has not been possible to create an in-silico approach that would reproduce the effects of immune tolerance.

Counter selection was introduced to the SELEX process to address this limitation. Counter selection works well at removing sequences that bind strongly to the counter selection target, but effectiveness decreases as cross reactivity decreases^19^. Furthermore, the weaker a given aptamer sequence cross-reacts to a counter selection target the lower the proportion of that aptamer that would be removed by the counter selection process^19^.

This is a fundamental problem given the magnitude of differences in the abundance of proteins in biological matrices and target proteins. For example, human serum albumin (HSA) is present in human blood at a concentration range of 522-746 μM^20^. If a target molecule was present at a concentration of 600 pM, then there would need to be a specificity of a million-fold towards the target over HSA, or the aptamer would be saturated by the weak binding to HSA over the aptamer. Counter selection against aptamer sequences that bind to HSA with an affinity that is 10,000 to 100,000-fold less strong than their affinity to the target molecule will not be effective in reducing the enrichment of such sequences in SELEX selection. We postulate that the binding of aptamers to abundant proteins in biological matrices has been the primary reason for the lack of commercial success of aptamers in therapeutics and diagnostics.

We have changed aptamer selection through a fundamental redesign which we have termed the Neomer library. This library design comprises sixteen random nucleotides interspersed with fixed sequences, thus reducing the number of possible sequences to 4.29 × 10^9^. This enables the application of the same set of sequences in selection against different targets or in replication against the same target. In this paper we describe the use of a Neomer library for the development of aptamers for interleukin 6 (IL6). We also provide a process for the in-silico evaluation of the performance of the selected IL6 sequences in a selection against HSA.

## Results

A comparison of the Neomer library design with the SELEX process is provided in Fig. 1A. This prediction was calculated by RNAFold v2.5.0 ^22^, using default parameters and a temperature of 22 °C. This lack of secondary structure is an important element of the library design. This means that any additional secondary structure will be driven by the identity of the random nucleotides.

**Figure 1:**
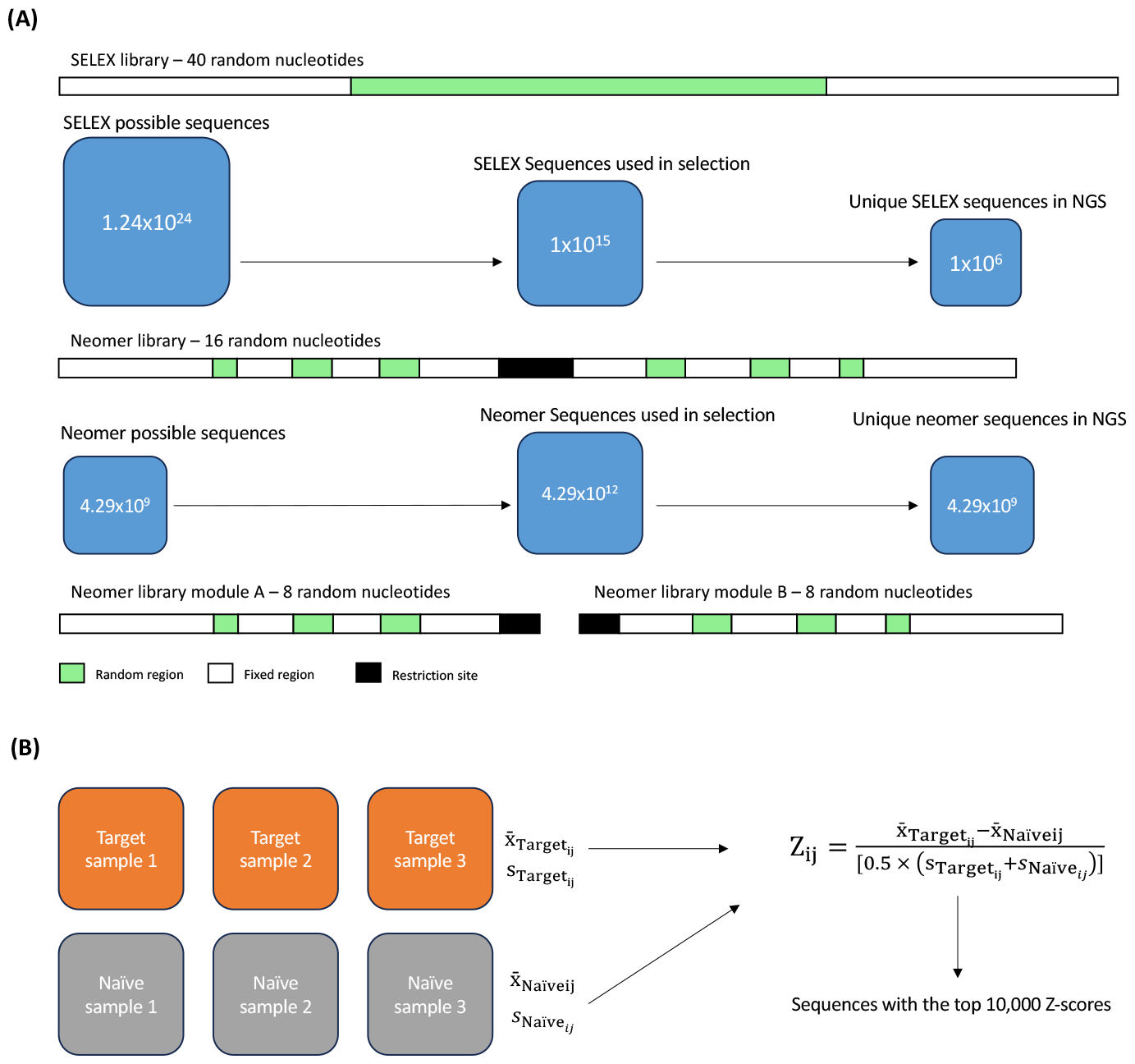
Visualization of the differences between SELEX and Neomer selection. **A)** A schematic showing the templates of SELEX and Neomer libraries respectively. A legend detailing the regions is provided at the bottom. Possible sequences, sequences used in selection and the number of unique sequences is shown in proportion below each template. **B)** A schematic showing how the z-score metric is evaluated in the Neomer approach.

To evaluate how the reduction of the total number of sequences in the Neomer library resulting from fewer random nucleotides impacts structural diversity, a comparison was conducted with the SELEX library. A reduction to sixteen contiguous random nucleotides in a SELEX library would significantly reduce the structural diversity of the library. The Neomer library has been designed such that this reduction in random nucleotides is possible without diminishing structural diversity. Specifically, the SELEX library design with a contiguous random region implicitly constrains the formation of secondary structure as adjacent nucleotides are limited in their capacity to fold. For example, if there is a sequence of ‘GGGCCC’, the secondary structure can be depicted using ‘……’ where ‘.’ represents an unhybridized nucleotide. The addition of any nucleotide, represented as ‘N’, would give us a sequence of ‘GGGNNNCCC’ and the character string changes to ‘(((…)))’ where ‘(‘ and ‘)’ represents the 5’ and 3’ sides of a hybridized region, respectively. It was observed that the contiguous nature of the first sequence is a constraint to the creation of secondary structure. This constraint is removed by the insertion of nucleotides between the bases with the potential to hybridize to each other.

Subsequently, the fixed sequences in the library were designed in to have minimum potential to hybridize with each other, ensuring secondary structure is driven by identity of the random sequences, either amongst themselves, or with the fixed sequences (Fig. 2).

**Figure 2:**
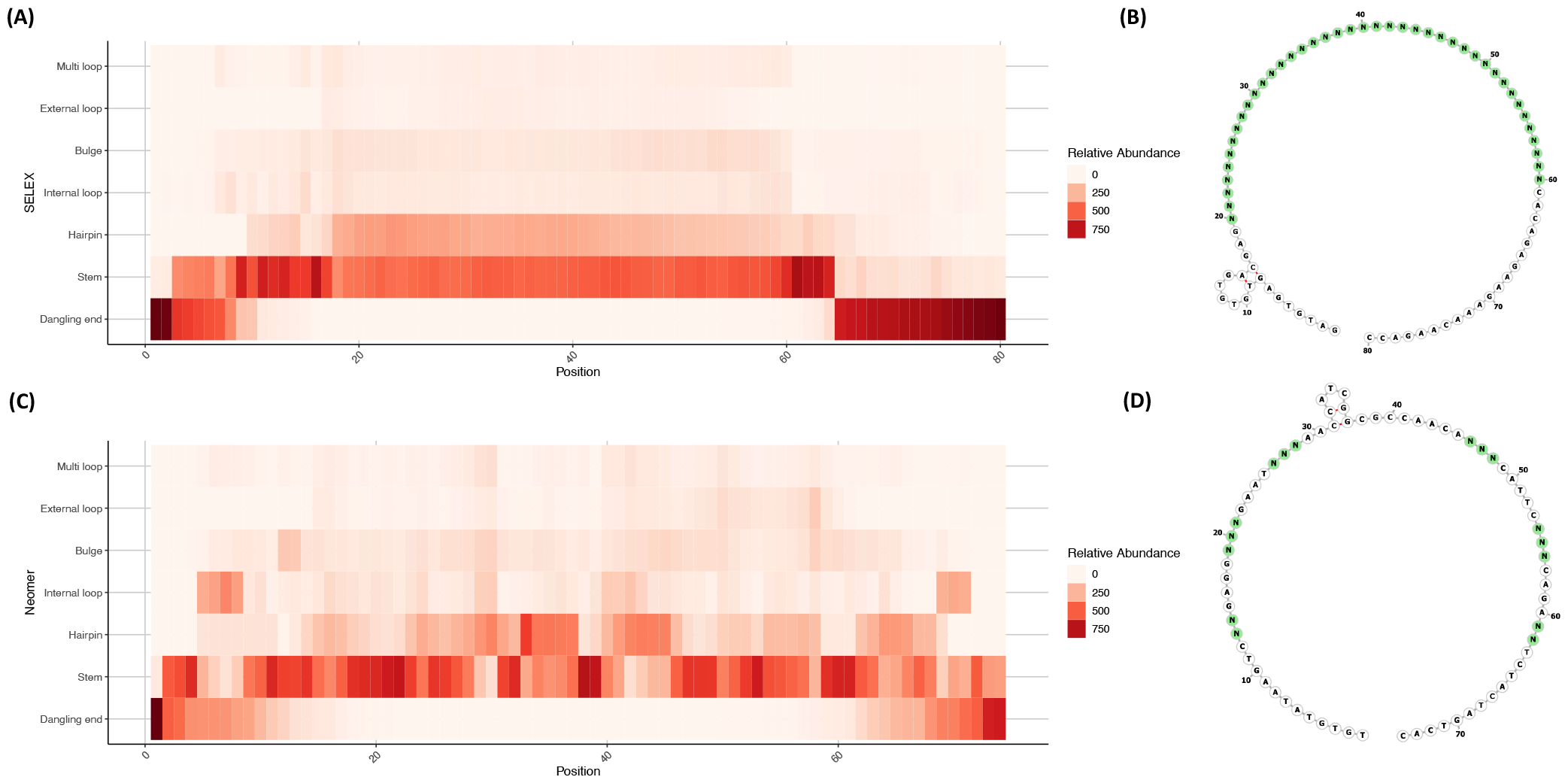
Comparison of the predicted secondary structures for SELEX and Neomer libraries. **A)** Heatmap plot of the distribution of secondary structure motifs across 1000 randomly generated SELEX template sequences. Relative abundance is detailed in the legend on the right. **B)** FORNA RNAfold plotting of the secondary structure of the SELEX template sequence. Random nucleotides are shown in green. **C)** Heatmap plot of the distribution of secondary structure motifs across 1000 randomly generated Neomer template sequences. Relative abundance is detailed in the legend on the right. **D)** FORNA RNAfold plotting of the secondary structure of the Neomer template sequence. Random nucleotides are shown in green.

To further compare structural diversity between the Neomer library and a SELEX library we randomly generated 1000 sequences using the Neomer template and a SELEX template that was previously published^21^. We computed the secondary structure of these sequences using Vienna RNAfold and annotated them using bpRNA^22,23^.

The Neomer and SELEX library differed in terms of secondary structure distribution, with the Neomer library showing a broader distribution of secondary structures and shorter dangling ends than the SELEX library (Fig. 2A and C). The probability that any given nucleotide would be contained within a specific structural motif was similar between the two library designs (Supp. Fig. 1). The position of these structures within the overall sequence was more broadly distributed in the Neomer library than in the SELEX library (Fig. 2). As such, the reduction of random nucleotides in the Neomer library does not imply a reduction in structural diversity compared to a SELEX library (Supp. Fig. 2).

The Neomer selection process starts with a pool of 4.29 × 10^12^ sequences, resulting in an average representation of 1,000 copies for each of the possible 4.29 × 10^9^ sequences. Moreover, by starting with an average of 1,000 copies of each sequence it is possible to assess the effect of selection on each sequence after a single round of selection, reducing the impact of PCR bias.

We cannot determine the frequency of each of the 4.29 × 10^9^ sequences with a single next generation sequencing (NGS) run. To mitigate this constraint, we divided each sequence into two parts (module A and B), each with eight random nucleotides or 65,536 possible sequences. NGS analysis was performed separately on each module of the library thus enabling sufficient NGS read coverage to predict the frequency of each possible sequence per module (Fig. 1B). The predicted frequency for each possible sequence within each module were multiplied by each other resulting in a predicted frequency for each of the 4.29 × 10^9^ sequences in the full possible sequence space.

A key feature of this library design is that it makes it possible to apply the same initial naÏve sequences to the same target multiple times, as well as apply the same initial naÏve sequences to different targets. The selection was performed with the same library on a target in triplicate, and on a negative control in triplicate (Fig. 1B).

We observed on average, 16 million reads for each module A and B replicate in NGS analysis (Supp. Table 1). A read count of over 5 million in each module indicated acceptable sequence coverage for each library analysed. A read count of 5 million represents an average of 76 copies per possible sequence. Frequencies for each corresponding sequence were complied across each replicate to calculate an average frequency and standard deviation of the average.

A Z-score metric was utilized to evaluate the performance for each of the 4.29 × 10^9^ possible sequences in selection, where the average predicted frequencies and standard deviation of these averages for both positive target (IL6) and negative controls (initial naÏve library) replicates were calculated. The top 10,000 sequences identified to have the highest z-scores were retained for further analysis (Fig. 1B).

Fig. 3A provides the distribution of fold values (IL-6/naÏve -1) versus the top 10,000 sequences based on Z-score values as calculated above. Sequences that exhibit both high fold and high z-score values would be expected to perform the best as candidate aptamers.

**Figure 3:**
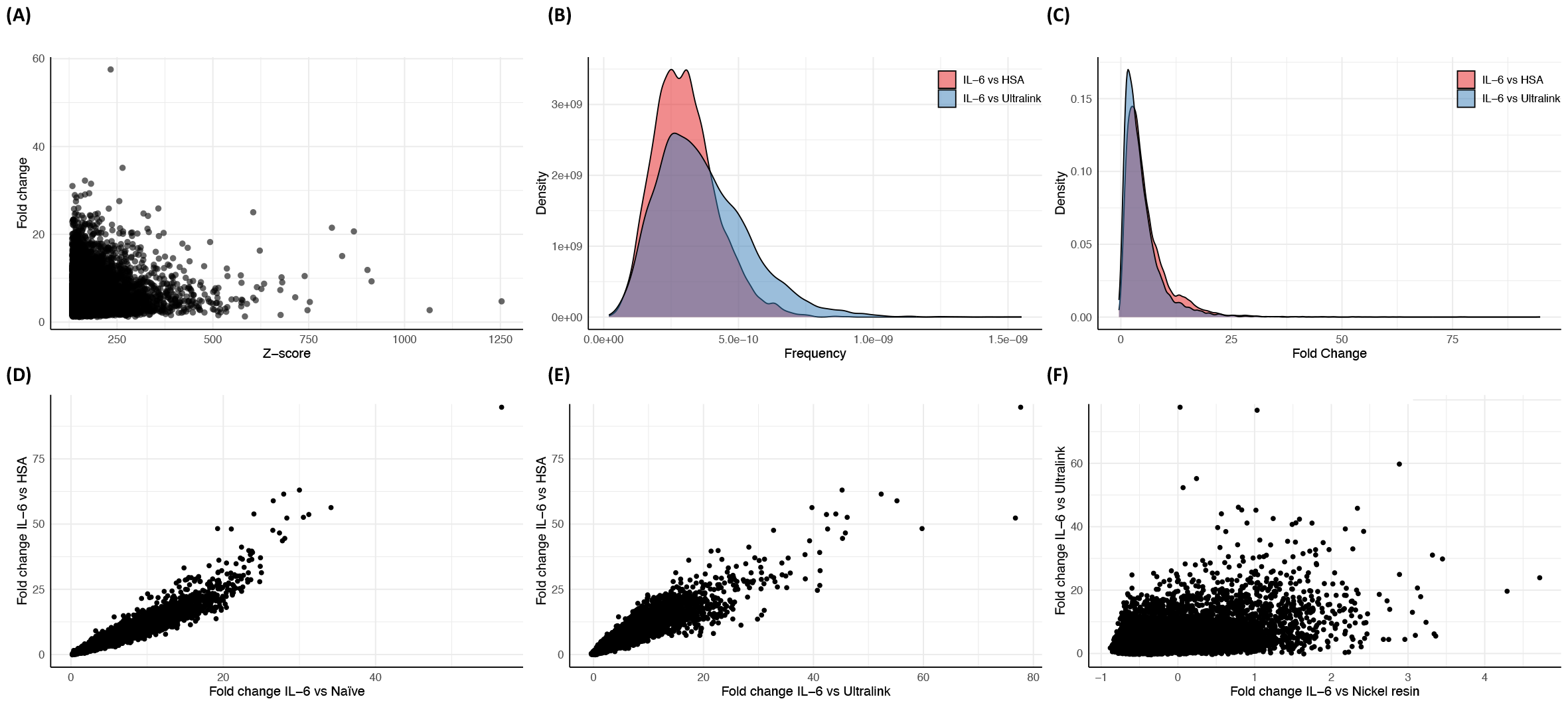
Analysis of the Next-Generation Sequencing results. **A)** Scatterplot of the Z-score vs fold change in the top 10,000 sequences identified from the IL-6/NaÏve selection. **B)** Densigram of the frequency of the top 10,000 IL-6/NaÏve selection sequences from the IL-6/HSA and IL-6/Ultralink selections. **C)** Densigram of the fold change of the top 10,000 IL-6/NaÏve selection sequences from IL-6/HSA and IL-6/Ultralink selections. **D)** Scatterplot of the fold change values from the top 10,000 IL-6/NaÏve selection sequences in the IL-6/NaÏve and IL-6/HSA selections. **E)** Scatterplot of the fold change values from the top 10,000 IL-6/NaÏve selection sequences in the IL-6/Ultralink and IL-6/HSA selections. **F)** Scatterplot of the fold change values from the top 10,000 IL-6/NaÏve selection sequences in the IL-6/Nickel resin and IL-6/Ultralink selections.

The use of the same 4.29 × 10^9^ sequences also enabled screening for the performance of these top 10,000 sequences on several counter targets. In this case, we screened for the performance of these sequences against a selection for HSA, against the UltraLink resin that the HSA was immobilized on, and against the nickel resin that the IL6 protein was immobilized on. The performance of these sequences on these counter targets was evaluated by comparing the frequency predicted on the positive target by the frequency predicted on each of these counter targets as a fold value (Fig. 3B, C, and D).

The overall distribution of fold differences between the IL6 sequences and counter-targets (HSA and UltraLink) was similar. This implies that sequences that exhibited high specificity against HSA also exhibited high specificity against the UltraLink resin.

Fig. 3D, E and F illustrate the level of concordance among sequences in specificity against various counter-targets. A high level of concordance between the fold values for the top 10,000 sequences in IL6 versus HSA and UltraLink was observed. This concordance was not apparent in comparison to the resin used for immobilization of IL 6, nickel resin. This indicates that certain sequences from the IL6 selection was selected based on their capacity to bind to the nickel resin rather than IL6. Multiple filters were required to identify the top candidate sequences in terms of specificity against all targets.

Table 1 shows the fold values for the top 15 IL6 sequences IL6 versus HSA fold values. The sequences in the table were sorted by IL6/HSA fold change. Four candidate sequences were further selected based on high performance and specificity against this group and nickel resin (IL-6-9805, IL-6-4202, IL-6-6449 and IL-6-7326.

**Table 1:**
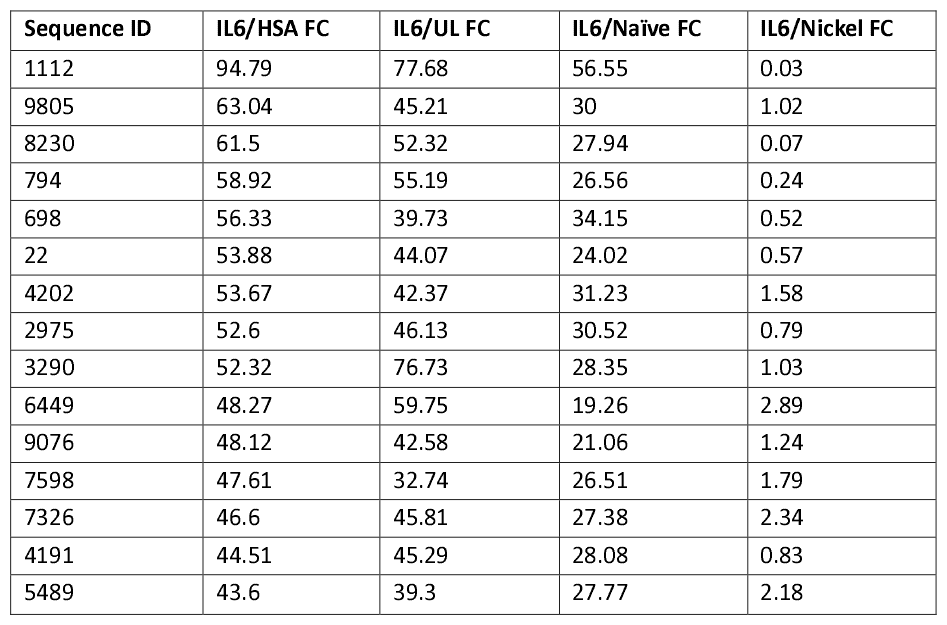
IL 6 Fold values for all contrasts.

The structures for these selected candidate sequences are presented in Fig. 4. Nucleotides highlighted in green represent the identity of random positions in the library. The identity of the random sequences in each case is provided in Table 2.

**Table 2:**
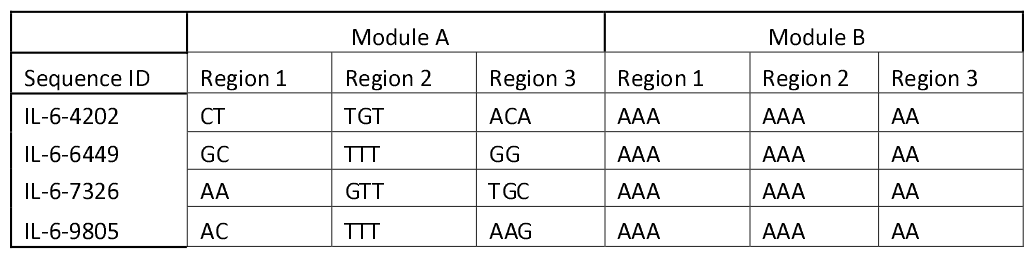
Identity of random module in IL-6 candidate sequences.

| Sequence ID | Module A |  |  | Module B |  |  |
| --- | --- | --- | --- | --- | --- | --- |
|  | Region 1 | Region 2 | Region 3 | Region 1 | Region 2 | Region 3 |
| IL-6-4202 | CT | TGT | ACA | AAA | AAA | AA |
| IL-6-6449 | GC | TTT | GG | AAA | AAA | AA |
| IL-6-7326 | AA | GTT | TGC | AAA | AAA | AA |
| IL-6-9805 | AC | TTT | AAG | AAA | AAA | AA |

**Figure 4:**
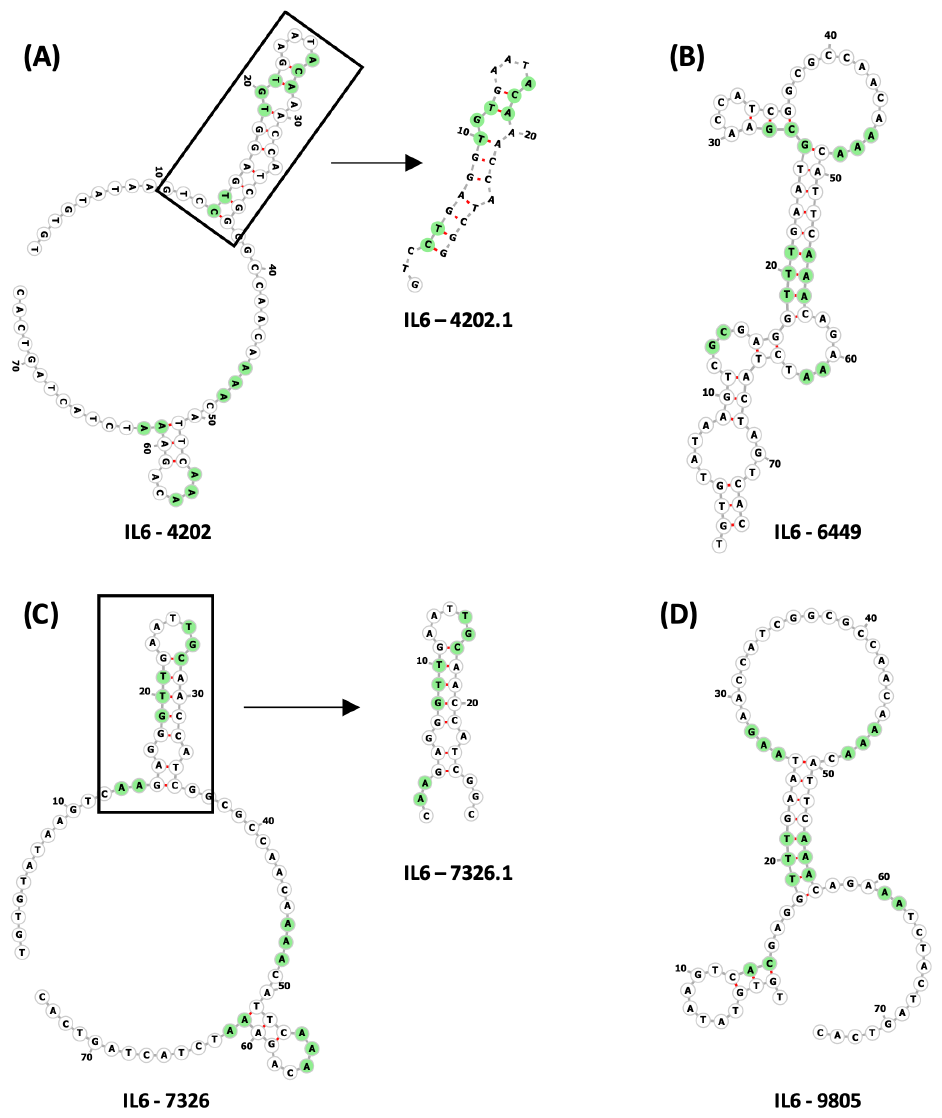
Depiction of the selected aptamer full-length and truncated sequences. **A)** FORNA visualisation of IL-6-4202 for plotting out secondary structure. Random nucleotide positions are labelled in green. A black box shows the truncated structure IL-6-4202.1. Secondary structure visualisation shows maintenance of the same structure from IL-6-4202. **B)** FORNA visualisation of IL-6-6449 for plotting out secondary structure. Random nucleotide positions are labelled in green. **C)** FORNA visualisation of IL-6-7326 for plotting out secondary structure. Random nucleotide positions are labelled in green. A black box shows the truncated structure IL-6-7326.1. Secondary structure visualisation shows maintenance of the same structure from IL-6-7326. **D)** FORNA visualisation of IL-6-9805 for plotting out secondary structure. Random nucleotide positions are labelled in green.

Prior to performing binding assays candidate sequences IL-6-4202 and IL-6-7326 were truncated with removal of a portion of the dangling ends. The truncations in both cases effectively removed module B from the sequence. All the top fifteen sequences listed in Table 1 were observed to contain the same B module sequence. High structural diversity was observed across candidate aptamers despite the nucleotide consistencies of the random regions in module B (Table 2). These nucleotides specifically contributed to varying structural motifs within the aptamers such as internal and external loops and stems (Fig. 4).

We injected varying concentrations of IL-6 and HSA in flow over the candidate aptamers as well as a negative control. The resonance due to binding is the result of the total resonance measured on the candidate aptamers minus the resonance measured on the negative control (Total resonance – flow through resonance = resonance due to binding). These results for each candidate aptamer are provided in Fig. 5. The calculated binding coefficients from this analysis are provided in Table 3. All four sequences exhibited strong binding affinity and high specificity to IL6. No binding was quantified when tested against HSA.

**Table 3:** Binding coefficients for candidate aptamers versus IL 6 and HSA.

| Sequence ID | IL 6 |  |  | HSA |
| --- | --- | --- | --- | --- |
| | Kd ($s^{-1}$) | ka ($M^{-1}s^{-1}$) | kD (M) | kD (M) |
| IL-6-4202.1 | 2.98E-03 | 4.07E+04 | 7.32E-8 | NB |
| IL-6-6449 | 1.52E-03 | 4.73E+04 | 3.21E-8 | NB |
| IL-6-7326.1 | 1.50E-03 | 5.56E+04 | 2.70E-08 | NB |
| IL-6-9805 | 1.77E-03 | 4.37E+04 | 4.05E-08 | NB |

NB = No binding measurable

**Figure 5:**
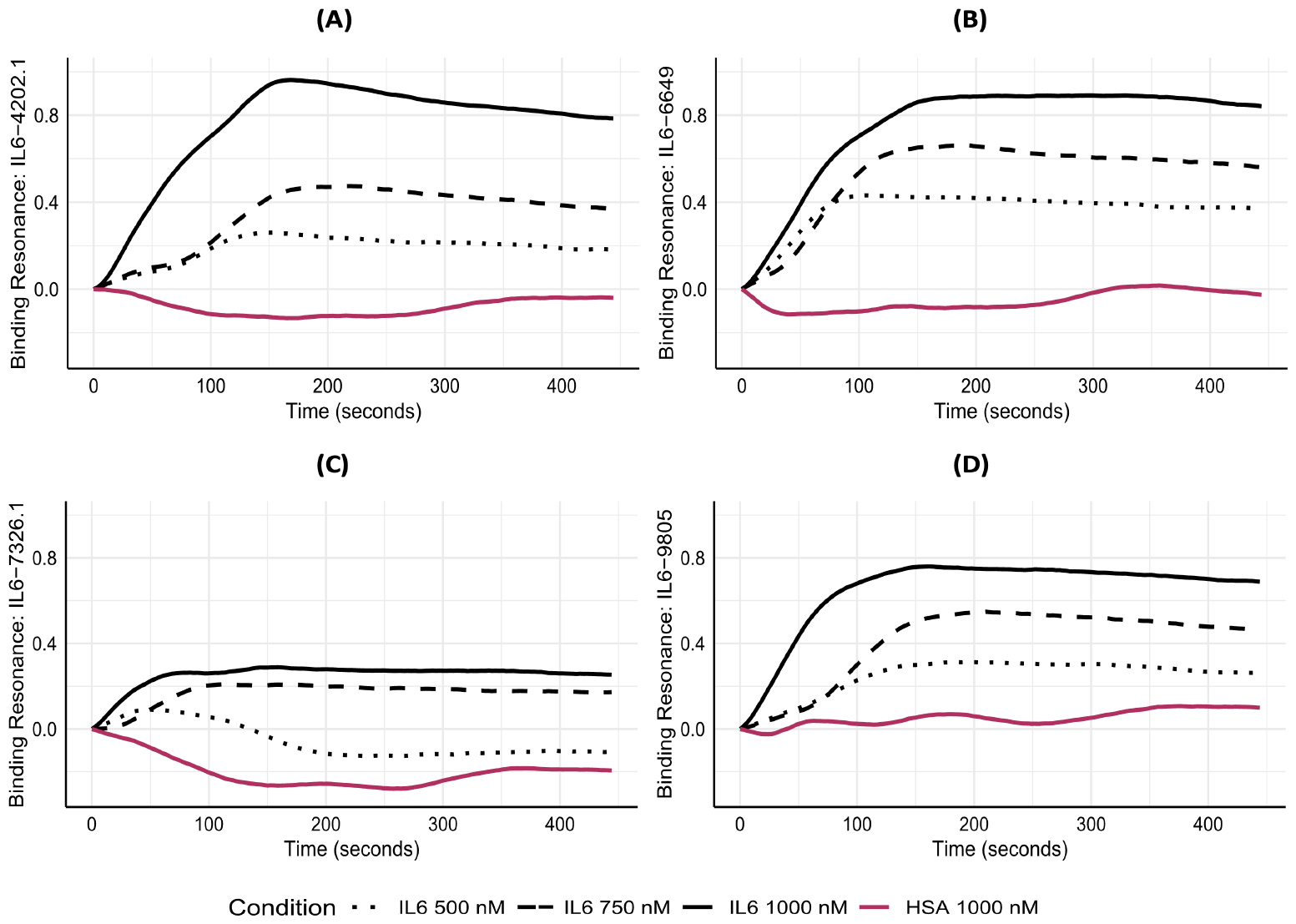
SPRi results for IL6 aptamer sequences to IL6 and HSA. **A)** Binding resonance of IL-6-4202.1 over time across a variety of concentrations of IL-6 detailed in the legend on the bottom. **B)** Binding resonance of IL-6-6649 over time across a variety of concentrations of IL-6 detailed in the legend on the bottom. **C)** Binding resonance of IL-6-7326.1 over time across a variety of concentrations of IL-6 detailed in the legend on the bottom. **D)** Binding resonance of IL-6-9805 over time across a variety of concentrations of IL-6 detailed in the legend on the bottom.

## Discussion

A novel approach to aptamer development has been demonstrated to effectively identify aptamers with sufficient binding affinities and specificity to be of interest for commercial application in diagnostics.

The Neomer library introduces an innovative design that is composed of sixteen random nucleotides interspersed with fixed sequences. The separation of the random nucleotides with fixed sequences that are designed to hybridize to each enables a maintenance of structural diversity when compared to 40 nucleotide contiguous random regions in a SELEX library. The reduction in the number of possible sequences in the Neomer library enables reproducible selection of the same sequences. In addition, by reducing the sequence solution space it is possible to initiate selection with multiple copies of each possible sequence, enabling a single round of selection.

Applying the Neomer method resulted in the successful identification of four aptamers with high selectivity and specificity for IL 6. The diagnostic application of these aptamers for the detection of IL 6 in plasma is being developed and will be detailed in the future.

The outer product calculation does not have the capacity to fully determine individual sequence frequencies. An artifact of this analysis method is the distribution of the enrichment of a given sequence among sequences that share a common module. The accuracy of sequence frequency prediction is affected by the magnitude of selection effect as a function of sequence enrichment. We have recognized that the addition of more replications of the naÏve control library reduces sampling error and decreases statistical artifacts due to low standard deviation values across a limited number of replications. This approach will be detailed in upcoming publications.

The creation of a closed sequence set enables reproducible selection of aptamers for the first time. Reproducible selection on the same target and on other targets provides a basis for robust statistical analysis of aptamer enrichment that was not previously possible with SELEX. Antibody selection is also not reproducible given host individual variation in pre-existing antibody repertoires and randomness introduced through V(D)J recombination^24^. The design of Neomer libraries thus provides an improvement on both SELEX and antibody development by enabling the addition of knowledge from multiple selections with the same initial naÏve repertoire.

The introduction of the Neomer approach signifies a pivotal advancement in aptamer development that will result in higher levels of specificity, overcoming a central bottleneck in aptamer commercialization for applications in diagnostics, drug delivery, and therapeutics. This innovation introduces the potential improvement of aptamer performance in diverse biological matrices, inciting a paradigm shift in the landscape of molecular recognition technologies.

## Materials and Methods

### Library Design

All DNA sequences including library, primers, and aptamers were synthesized by Integrated DNA Technologies with standard desalting. The Neomer library was designed as synthetic ssDNA, 73 nt long with 16 total randomized nt interspersed between fixed sequences. The fixed sequences include flanking primer binding sites, and a KasI restriction site in the middle (5’-TGT GTA TAA GTC NNG AGG NNN GAA TNN NAA CCA TCG GCG CCA ACA NNN CAT TCN NNC AGA NNT CTA CTA GTC AC 3’; where ‘N’ represents an equal probability of any nucleotide).

### Target preparation

#### Interleukin 6 (IL6)

Targets for selection were immobilized on resin scaffold. Recombinant IL-6 with an N-terminal his tag (Fitzgerald, Biosynth Ltd.) was immobilized on His-Pur™ Ni-NTA Resin (ThermoScientific) following a modified purification protocol. Il6 (50 μg) was prepared in 200 μL of binding buffer (20 mM sodium phosphate, 300 mM sodium chloride, pH 7.4) and incubated with 70 μL of Ni-NTA Resin for 2 hr at room temperature with agitation. Unbound protein was collected by centrifuging at 700 x g for 2 min and removing the supernatant. The resin was washed once with 140 μL of binding buffer. Remaining active sites were blocked with 2 mM imidazole, incubating for 1 hr at room temperature with rotation. After blocking resin was washed with 1x PBS and resuspended as a 50:50 slurry in 1x PBS.

#### Human Serum Albumin (HSA)

HSA (Sigma-Aldrich Canada) was immobilized on UltraLink Biosupport (ThermoScientific) following user guide recommendations. Solubilized HSA was prepared to a final concentration of 2 mg/mL in conjugation buffer (500 mM sodium citrate, 200 mM sodium bicarbonate, pH 8.5). Added 200 μL of protein solution to 8.8 mg of dry UltraLink Beads (swell volume of 8 μL/mg) and incubated for 2 hr at room temperature with agitation. Unbound protein was collected by centrifuging at 1200 x g for 5 min. The resin was washed once with 140 μL of conjugation buffer. Remaining active sites were blocked overnight with 1 M Tris at 4 °C. After blocking the resin was washed and stored in PBS.

### Neomer Selection

Prior to selection, the Neomer library was denatured at 95 °C for 5 min, cooled on ice for 10 min, and equilibrated to room temperature. Selection was conducted in a 1 mL column fitted with a 20 μm frit. The refolded library (7.15 pmoles) was incubated with 10 μL of IL6-Ni-NTA resin (223 pmol IL6) in 100 μL of selection buffer (10 mM HEPES, 120 mM sodium chloride, 5 mM potassium chloride, 5 mM magnesium chloride, pH 7.6) for 30 min at room temperature with agitation. The unbound library was discarded in the flowthrough and the column washed three times with 500 μL of selection buffer. Bound library was eluted by adding 200 μL of 6 M urea to the column, heating at 85 °C for 5min and collecting the flowthrough. The elution was repeated, and the eluents pooled together. The eluted library was purified using the GeneJet PCR Purification Kit (ThermoFisher) and eluted with 50 μL of MilliQ filtered water. The library was brought up to 100 μL in selection buffer and incubated with another 10 μL of IL6-Ni-NTA resin. After the second selection, the purified library was eluted in 400 μL of MilliQ filtered water and stored in a salinized vial. Selection against HSA was conducted using the same procedure with the following changes. The library was incubated with 10 μL of HSA-UltraLink Biosupport (369 pmol HSA) in 100 μL of PBS. To increase the yield of recovered library, the second selection column was omitted. The purified library was eluted in 400 μL of MilliQ filtered water. Each selection as described was conducted in triplicate for each target.

### NGS Preparation

Nested PCR primers were applied to each selected DNA library for sequence identification. The sequences were amplified (Supp. Table 2), isolated from a 20% acrylamide gel, and purified for sequencing. For purification of the DNA, NGS2 PCR products were run on 20% polyacrylamide gels at 150V for 5 hr. The target band was excised from the gel, fragmented, and stored in TE buffer in a salinized vial for 3 days to elute the DNA. The DNA was subsequently purified using a Genejet PCR purification kit. The DNA was purified following standard protocol and eluted from the column with 30 μl of water. 10 μl of the DNA is run on a 10% polyacrylamide gel to determine concentration of each library. Sequence analysis was completed by the Hospital for Sick Children (Toronto, CA) using Illumina HiSeq 2500.

Negative control selections of the library against UltraLink resin and nickel resin without immobilized proteins was also performed in triplicate as well as the unselected naÏve library.

### Z-score calculations

Next-generation sequencing of all libraries was performed by The Centre for Applied Genomics, The Hospital for Sick Children, Toronto, Canada. FASTQ files obtained upon sequencing were converted into a FASTA format, and the copy number of each of the possible 65,536 sequences in each of Module A and Module B were characterized using a proprietary Python script that we have developed for this application. The frequency of each Module sequence was determined by dividing copy number by the total number of reads for that Module as seen in equation (1). The frequency of each of the 4.29 × 10^9^ sequences in the original selected library was then estimated through an outer product calculation of the Module A and Module B frequencies.

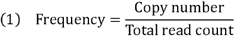

The average frequency for each of these 4.29 × 10^9^ sequences was determined for the three target samples and the three counter target samples in each selection. The standard deviation of each average was also determined. The average frequency of the target samples for each sequence was subtracted from the average frequency for the same sequence across the counter-target samples. This value was divided by the average of the standard deviation of these averages from both sets of samples to generate a Z score as seen in equation (2). The top 10,000 sequences in terms of Z score were identified.

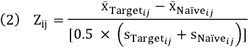

The perform.ance of each of these top 10,000 sequences was evaluated in selections against the counter targets: HSA, UltraLink resin, and nickel resin using fold analysis. The average frequency of each of these sequences was predicted for each counter target and divided into the average frequency predicted for that sequence on the positive target resulting in a fold value as seen in equation (3).

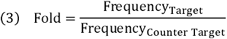

Where Target is the average frequency of each of the top 10,000 predicted aptamers for IL-6, and Counter Target is the corresponding average frequency of the same sequence in each of full counter target’s full selection. The data and script used to generate the data can be found at: https://github.com/Chloemansour/neomer_paper_pipeline

### Secondary structure analysis

1000 random sequences were generated using the Neomer and SELEX templates respectively^21^. Dotbracket structures for these were calculated using RNAfold 2.5.0 with parameters specifying a temperature of 22 °C and a DNA folding specific energy profile^22^. Secondary structure annotation for these sequences was computed using bpRNA under default parameters^23^. The data and script used to generate the data can be found at: https://github.com/Chloemansour/neomer_paper_structural_diversity.

### Visualisation of IL-6 aptamer analysis and secondary structure analysis

Data were processed in python 3.9.13 and were then loaded into R studio (version 2023.09.1+494) where they were analysed and visualised using ggplot and other packages^25–28^. The data and script used to generate the data can be found at: https://github.com/Chloemansour/neomer_paper_visualizations.

### Aptamer library template visualisation

Secondary structure dotbracket information was computed using RNAfold 2.5.0^22^ with parameters specifying a temperature of 22 °C and a DNA folding specific energy profile and then inputted into FORNA^29^. A custom colour palette was chosen to highlight random regions in each library. The data and script used to generate the data can be found at: https://github.com/Chloemansour/neomer_paper_visualizations.

### General SPRI Materials and Methods

Binding assays were performed using the SPRi system (OpenPlex, HORIBA France). Bare gold biochips were purchased from Xantac Bioanalytics (Düsseldorf, Germany). All thiolated aptamers were synthesized and purchased from Integrated DNA Technology (IDT) (Lowa, USA). Recombinant Human IL-6 protein was purchased from Peprotech (Cranbury, USA) Albumin from human serum (HSA) was purchased from Sigma-Aldrich Canada. The 1x Running Buffer (pH5.7±0.1 at 25 °C) contains 400 mM 6-aminohexanoic acid (EACA) (Sigma-Aldrich Canada), 2 mM Triethylenetetramine (TETA) (Sigma-Aldrich Canada) and 100 mM NaCl (Sigma-Aldrich Canada). The blocking agent 6-Mercapto-1-hexanol and Phosphate-Buffered Saline (PBS) (Sigma-Aldrich Canada) (10.14 mM Na_2_PO_4_, 137 mM NaCl, 2.68 mM KCl, pH 7.4) at 25 °C. The calibration solution in 1x running buffer contains 2mg/mL Sucrose (Sigma-Aldrich Canada) and the regeneration solution for gold chip is 1%(w/v) Sodium Dodecyl Sulfate (SDS) (Sigma-Aldrich Canada).

### Preparation of Sensor Chip

The sensor chip and purified salt free thiolated aptamers were brought to room temperature prior to spotting, aptamers were immobilized on the gold surface in triplicate at 10 nL/spot with a concentration of 5 μM. The gold chip was allowed a 1-hour incubation time for efficient immobilization at room temperature and at an approximate humidity of 80%. 1mM 6-mercapto-1-hexanol in 1x PBS was used to facilitate blocking of the functionalized surface of gold chip for an hour. The prepared sensor chip was then placed into the instrument.

### SPRi Measurement

Surface plasmon resonance can be explained by an optical detection process that happens when a gold-layered prism is hit by a polarized light which facilitates the conversion from light photons into surface plasmon waves. This allows the measurement of a change of resonance condition due to any interaction between immobilized probes and targets. OpenPlex SPRi systems allows real-time quantification and monitoring of the biomolecular interactions. The method is to calculate the angle where the greatest variation of reflectivity Δ%R will be and adjusting this angle to monitor kinetics activity which is the amount of reflectivity versus time. A CCD camera is used in this system to visualize the spots and monitor their reflectivity simultaneously.

The biochip was loaded into the flow cell and target prepared in 1x running buffer was injected into the 200 μL sample loop at a flow rate of 50 μL/min. One completed run is 9 minutes, this means that the target was in flow over the aptamers for 240 seconds which is considered as association phase (ka). The remaining 300 seconds which is disassociation phase (kd) indicates the target was no longer in flow. Equilibrium dissociation constant (kD) was calculated using the equation kD = kd/ka. SPRi response recorded was the average response of 3 spots of each immobilized aptamer, the non-specific (negative control) signal was subtracted during data analysis. A calibration coefficient was obtained by injecting 2 mg/mL sucrose in running buffer to collect a variation of the reflectivity of individual spot which used to fix all spots to the same change in reflectivity, this promoted a kinetics camera angle of 58.16 degree. 1% (w/v) SDS solution was utilized to regenerate the functionalized surface, removal efficiency was measured by SPR signals, once the signal was restored to baseline the sensor chip was ready for another target injection. All injections were conducted at 25 °C (room temperature).

## Supporting information

Supplementary Figues and Tables

## Contributions

Conception and design of library: G.P. Optimization of library selection and NGS processing approaches, G.P., E.L.H., and S.L. Acquisition, analysis or interpretation of data: G.P., C.M., C.G.M., C.J.D. and Y.A. Writing and/or revisions of the manuscript: G.P., C.M., C.G.M., and E.R. All authors have approved the submitted version of the manuscript; agree to be personally accountable for their own contribution; and commit to any action that ensures or assesses integrity of any part of the work.

## Competing interests

C.M. and S.L are employees of NeoVentures Biotechnology Europe SAS, a privately held company that applies aptamer libraries to identify biomarkers. E.L.H., C.G.M., C.J.D., Y.A, and E.R. are employees are of NeoVentures Biotechnology Inc., a privately held company that develops aptamers for defined targets. G.P. is an owner of NeoVentures Biotechnology Inc. which in turn wholly owns NeoVentures Biotechnology Europe SAS.

