## Supplementary Figues and Tables for "A new method for the reproducible development of aptamers (Neomers)"

**Supp. Fig. 1: The probability that any given nucleotide would be contained within a specific structural motif was similar between SELEX and Neomer library designs.** **A)** Average **s**econdary structure motif frequency in each sequence across 1000 randomly generated SELEX sequences. **B)** Average contiguous nucleotide length of secondary structure motifs in each sequence across 1000 randomly generated SELEX sequences. **C)** Average **s**econdary structure motif frequency in each sequence across 1000 randomly generated Neomer sequences. **D)** Average contiguous nucleotide length of secondary structure motifs in each sequence across 1000 randomly generated Neomer sequences.

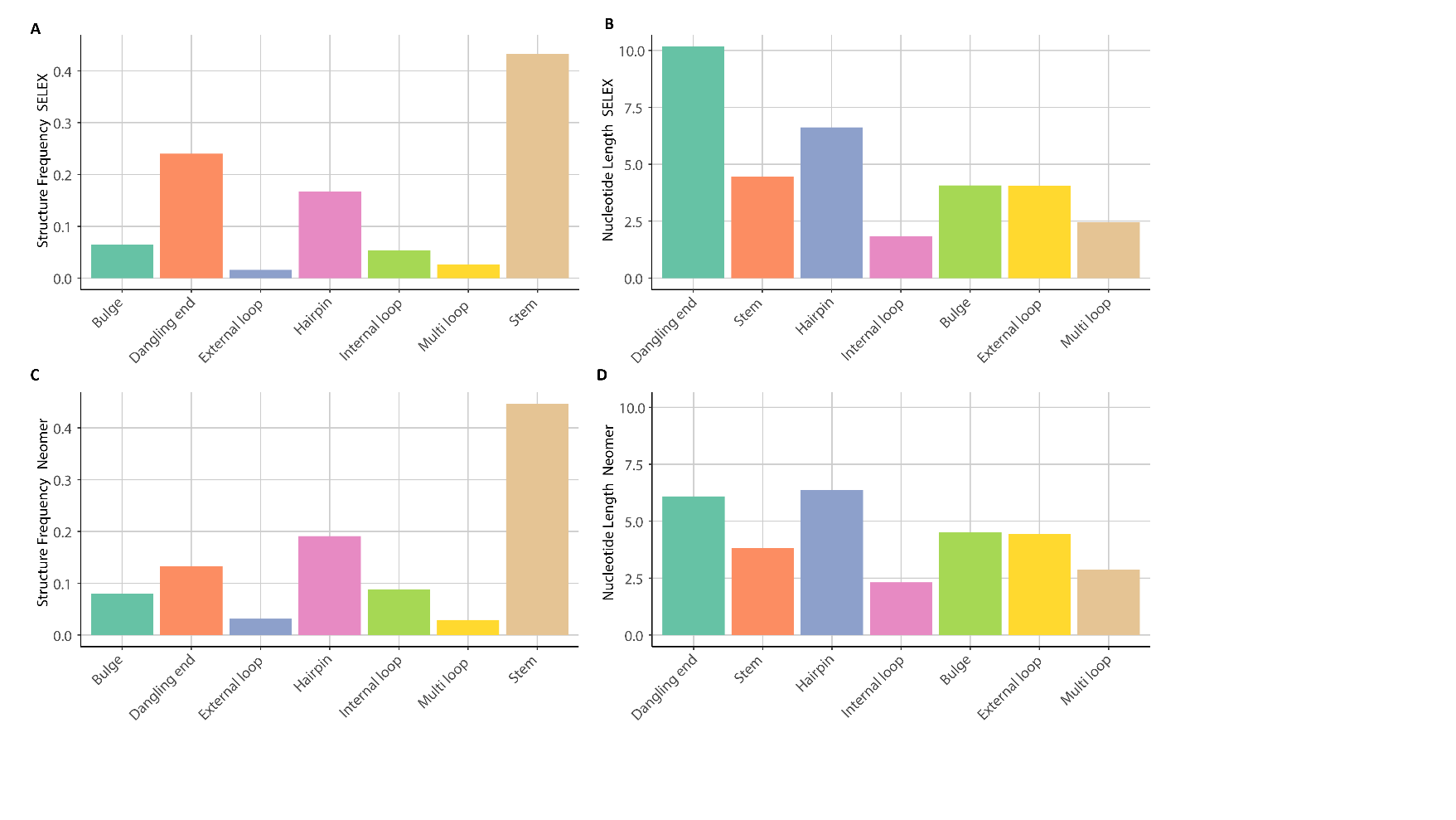

**Supp. Fig. 2: Average Shannon diversity index value across position for the SELEX and Neomer template. The template per position is plotted out below the plot.**

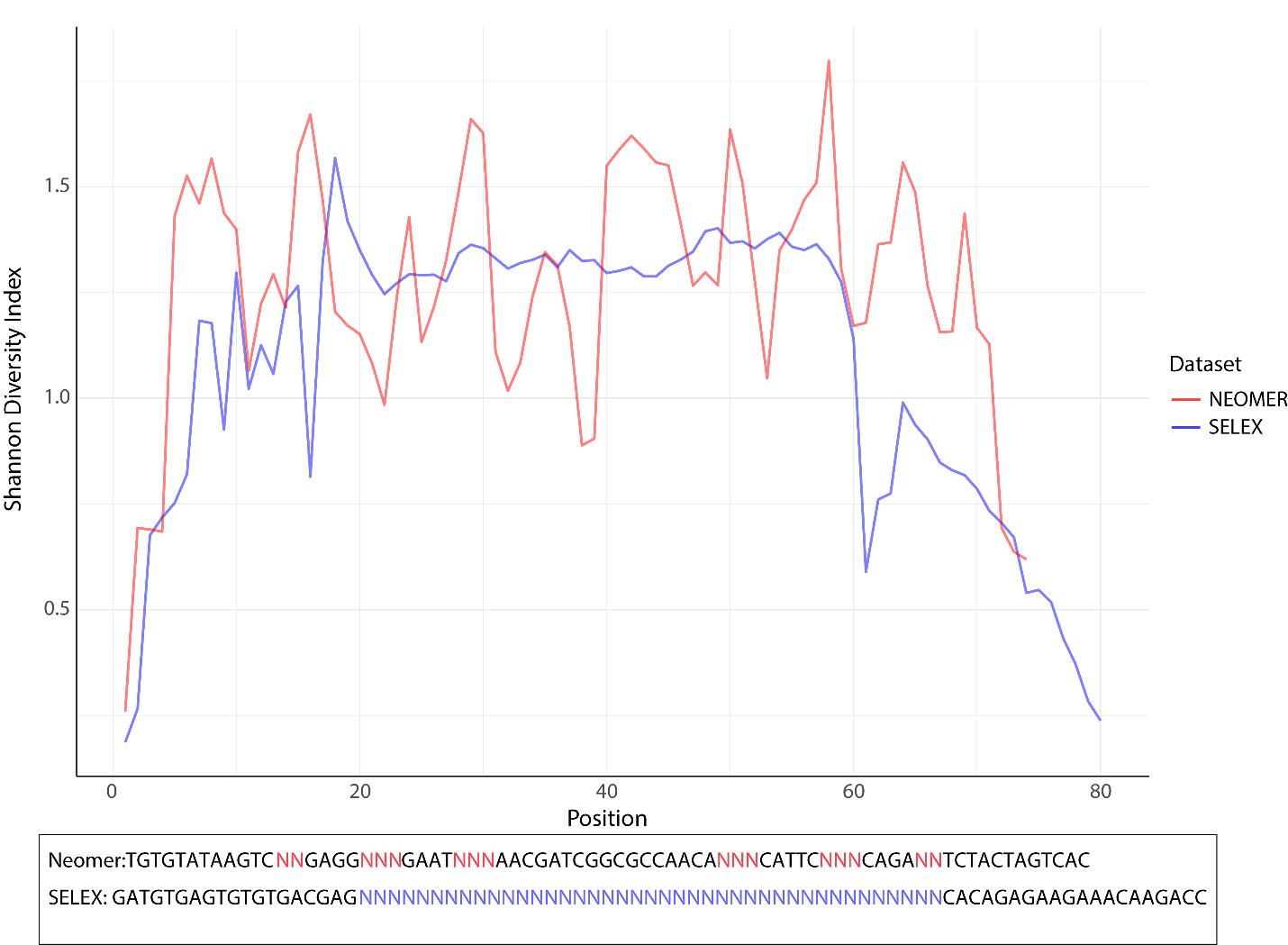

**Supp. Table 1: Reads from NGS.**

| **Sample Name** |  | **Module A Counts** |  | **Module B Counts** |  | **Total A + B Counts** |
| --- | --- | --- | --- | --- | --- | --- |
| IL 6 |  | 15,636,440 |  | 18,418,205 |  | 34,054,645 |
|  |  | 18,307,668 |  | 16,278,830 |  | 34,586,498 |
|  |  | 26,170,063 |  | 15,742,736 |  | 41,912,799 |
|  | Subtotal A: | 60,114,171 | Subtotal B: | 50,439,771 |  |  |
| HSA |  | 8,024,915 |  | 7,409,372 |  | 15,434,287 |
|  |  | 10,732,965 |  | 6,606,378 |  | 17,339,343 |
|  |  | 7,396,384 |  | 7,020,238 |  | 14,416,622 |
|  | Subtotal A: | 26,154,264 | Subtotal B: | 21,035,988 |  |  |
| Naive |  | 52,978,828 |  | 11,010,993 |  | 63,989,821 |
|  |  | 6,797,916 |  | 14,361,657 |  | 21,159,573 |
|  |  | 8,846,060 |  | 19,955,954 |  | 28,802,014 |
|  | Subtotal A: | 68,622,804 | Subtotal B: | 45,328,604 |  |  |
| Nickel Resin |  | 24,152,760 |  | 21,664,127 |  | 45,816,887 |
|  |  | 28,449,715 |  | 26,463,258 |  | 54,912,973 |
|  |  | 26,912,596 |  | 13,828,893 |  | 40,741,489 |
|  | Subtotal A: | 79,515,071 | Subtotal B: | 61,956,278 |  |  |
| UltraLink |  | 13,246,353 |  | 19,423,621 |  | 32,669,974 |
|  |  | 18,885,609 |  | 22,942,733 |  | 41,828,342 |
|  |  | 48,388,049 |  | 47,107,832 |  | 95,495,881 |
|  | Subtotal A: | 80,520,011 | Subtotal B: | 89,474,186 |  |  |
|  | **Total A counts:** | 314,926,321 | **Total B counts:** | 268,234,827 | **Grand total:** | 583,161,148 |

**Supp. Table 2: Primers used in NGS preparation.**

| Neoer III Fwd | CAAATACGTATGAGGTCGCTCGTTCTGTGTATAAGTC |
| --- | --- |
| Neoer III Rvs | TAATACGACTCACTATAGGGATAATGTGACTAGTAGA |
| RNm NGS1-A Fwd | CCCTACACGACGCTCTTCCGATCTNNNNNNCAAATACGTATGAGGTCGCTCGTTC* |
| RNm NGS1-A Rvs | GGTCAGACGTGTGCTCTTCCGATCGGGGCGCCGATGGTT |
| RNm NGS1-B Fwd | CCCTACACGACGCTCTTCCGATCTATCACGGCGCCAACA |
| RNm NGS1-B Rvs | GGTCAGACGTGTGCTCTTCCGATCGGGTAATACGACTCACTATAGGGATAATGCTGTCTACTG |
| NGS2 Fwd | AAT GAT ACG GCG ACC ACC GAG ATC TAC ACT CTT TCC CTA CAC GAC GCT CTT CCG |
| NGS2 Rvs | CAA GCA GAA GAC GGC ATA CGA GAT GTG ACT GGA GTT CAG ACG TGT GCT CTT CC |

*Where N represents the position of the Hex Code used for NGS data analysis
